# pcaReduce: Hierarchical Clustering of Single Cell Transcriptional Profiles

**DOI:** 10.1101/026385

**Authors:** Justina Žurauskienė, Christopher Yau

## Abstract

**Motivation:** Advances in single cell genomics provides a way of routinely generating transcriptomics data at the single cell level. A frequent requirement of single cell expression experiments is the identification of novel patterns of heterogeneity across single cells that might explain complex cellular states or tissue composition. To date, classical statistical analysis tools have being routinely applied to single cell data, but there is considerable scope for the development of novel statistical approaches that are better adapted to the challenges of inferring cellular hierarchies.

**Results:** Here, we present a novel integration of principal components analysis and hierarchical clustering to create a framework for characterising cell state identity. Our methodology uses agglomerative clustering to generate a cell state hierarchy where each cluster branch is associated with a principal component of variation that can be used to differentiate two cellular states. We demonstrate that using real single cell datasets this approach allows for consistent clustering of single cell transcriptional profiles across multiple scales of interpretation.

**Availability:** R implementation of *pcaReduce* algorithm is available from https://github.com/JustinaZ/pcaReduce

## 1 Introduction

Recent advances in high-throughput single cell RNA sequencing (RNA-seq) technology has provided a way of collecting quantitative gene expression measurements across large numbers of cells, tissue types and diversity of cellular states (Saliba *et al*., 2014; Macaulay and Voet, 2014; Shaleki *et al*., 2014; Sandberg, 2014; Kalisky and Quake, 2011; Wang and Bodovitz, 2010). For this reason single cell RNA-seq has increasingly became a method of preference in discovering molecular underpinnings of complex and rare cell populations (Achim *et al*., 2015; Buettner *et al*., 2015), assessing tissue composition (Zeisel *et al*., 2015; Jaitin *et al*., 2014; Scialdone *et al*., 2015), studying various diseases (Patel *et al*., 2014) and cell development/lineage processes (Satija *et al*., 2015; Deng *et al*., 2014; Marco *et al*., 2014; Treutlein *et al*., 2014).

A natural problem that has arisen from the availability of this technology is the development of analytical tools for the analysis and identification of cell sub-populations, cellular types or states from a single cell transcriptomics data (Stegle *et al*., 2015). Based on similarities and biological features, various computational techniques can assist not only with cell grouping problems but also with complex tasks, e.g. the determination of a number of clusters that describes the data in an optimal way. This can be either an entirely unsupervised operation, where no prior knowledge of clusters are assumed, or (semi-)supervised, where the number of clusters and their features maybe (or at least partially) known. Furthermore, it might be desirable to establish the relationships between the clusters and derive a cellular hierarchy that can capture both broad and specialised cellular classes.

For example, Figure 1 illustrates three clustering structures derived from a single cell study of mouse sensory neurons (Usoskin *et al*., 2015). Four broad sensory neuronal cell types (NF, TH, PEP, NP) were established by examining clusters of cells in the subspace spanned by the first few principal components (Figure 1A) and the differential expression of key cell markers. These four classes were then further sub-divided by iterative application of principal component analysis, using information contained in lower-ranked principal components (Figure 1B,C). Our proposed algorithmic contribution formalises such an approach. However, whereas this analysis was semi-supervised in that prior knowledge of marker genes was used to reliably identify the main four neuronal cell types. The question we asked was whether we could recapitulate such results fully automatically in an unsupervised fashion without any prior knowledge of cell type markers? This question is important in study conditions where there maybe little or unreliable prior knowledge of cell states/types.

**Figure 1:**
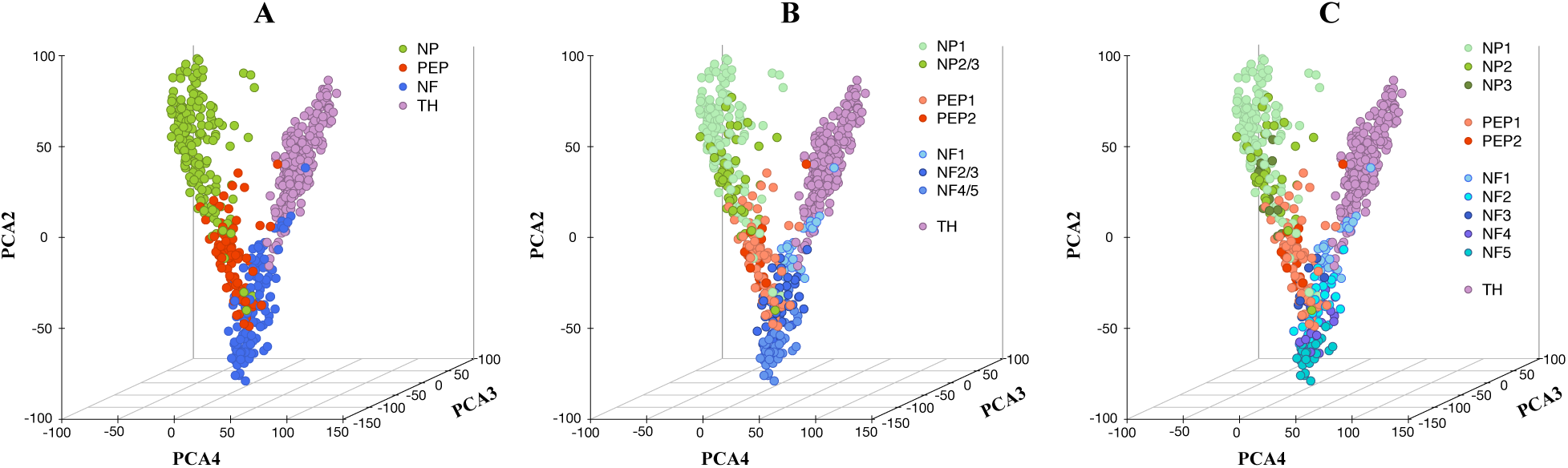
Cellular hierarchies. Three hierarchically related clustering structures for a single cell mouse neuronal dataset (Usoskin *et al*., 2015). The data has been projected on to the first four principle directions, we report the three that allows best data visualisation; we used the given cellular labels to colour cells according to the (A) 4, (B) 8, and (C) 11 cell subtypes identified in the original study.

A large number of clustering methods have been developed that can be applied to expression profile clustering (Jiang et al., 2004; Fraley and Raftery, 2004), this includes single-cell specific tools (Xu and Su, 2015) that are based on a combination of methodological approaches. Despite the catalogue of specialist methods available, generic techniques, e.g. hierarchical clustering, remain a popular choice; this is often due to their simplicity, flexibility, speed and surprising robustness in real-world analysis situations. Agglomerative hierarchical clustering treats each object initially as a singleton cluster and then successively merges (or agglomerates) pairs of clusters until all clusters have been joined into a single cluster, which contains all objects. This means that no explicit specification of the number of clusters is required *a priori*. A measure of dissimilarity between sets of observations is required to decide which clusters should be combined. This is typically achieved through the use of an appropriate metric (a measure of distance between pairs of objects), and a linkage criterion which specifies the dissimilarity of sets as a function of the pairwise distances of objects in the sets.

Here we have developed an agglomerative clustering approach that integrates principal components analysis (PCA) that we call *pcaReduce*. We first describe our method and then we detail the motivation behind this particular algorithmic design. Precisely, we show that our design corresponds to a coupling the reduced dimension data representation of each hidden layer of an linear autoencoder network to a corresponding layer in a hierarchical clustering structure. This interpretation provides a framework for future extensions of our tool. We demonstrate the effectiveness of *pcaReduce* for capturing cellular hierarchies using real single cell datasets and show that it performs as well, if not better, than other state-of-the-art approaches in an unsupervised setting.

## 2 Approach

### Algorithmic Overview

Let **X**_*n×d*_ denotes a gene expression matrix, where *n* is the number of cells measured across *d* number of genes; i.e. each cell **x**_*i*_ = {*x*_*i*1_, …, x_id_}is a *d*-dimensional object. Further assume that *X*_*ni*_ denotes a subset of *n*_*i*_ cells, *X*_*ni*_ ⊂ **X**. Our clustering algorithm begins by performing *K*-means clustering operation on the projection of the original gene expression matrix, **X***_n×d_*, to the top *K−*1 principal directions. The number of initial clusters *K* is set to a large value, say 30. Once the initial clustering is determined, we take two subsets (*X*_*ni*_, *X*_*nj*_) that originate from a pair of clusters (*i, j*) respectively, and calculate the probability for those observations to be merged together, *p*({*X*_*ni*_, *X*_*nj*_}μ_*ij*_, ∑_*ij*_). We assume that the probability density function is multivariate Gaussian with mean and covariance matrix given by:

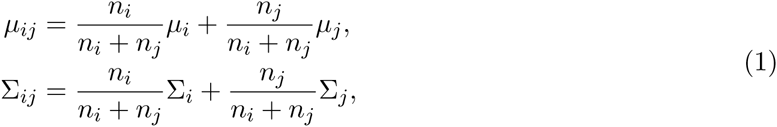

where (*n*_*i*_, *n*_*j*_), (*μ_i_, μ_j_*) and (Σ*_i_,Σ_j_*) denote the sizes, centroids and covariances of the clusters *i* and *j* respectively. We repeat this for all possible pairs (*i, j*). We then choose to merge two clusters by either (i) picking the pair that has the highest probability or (ii) sampling a pair of clusters to merge in proportion to their (normalised) merged probabilities. The number of clusters will now decrease to *K−*1. We now project the data matrix on to the first *K−*2 principal directions, i.e. removing the (*K−*1)-th principal component that explains the lowest degree of variance in the data, removing this dimension from the existing cluster centroids and covariance matrices. The above clustering operation is then repeated so that after every merge operation we remove a principal direction until only a single cluster remains. If sampling-based merge operations are used, the whole process can be repeated to obtain a number of alternative clusterings. This will be useful for assessing the stability of the clustering results. Algorithm 1 gives a pseudo-code description.

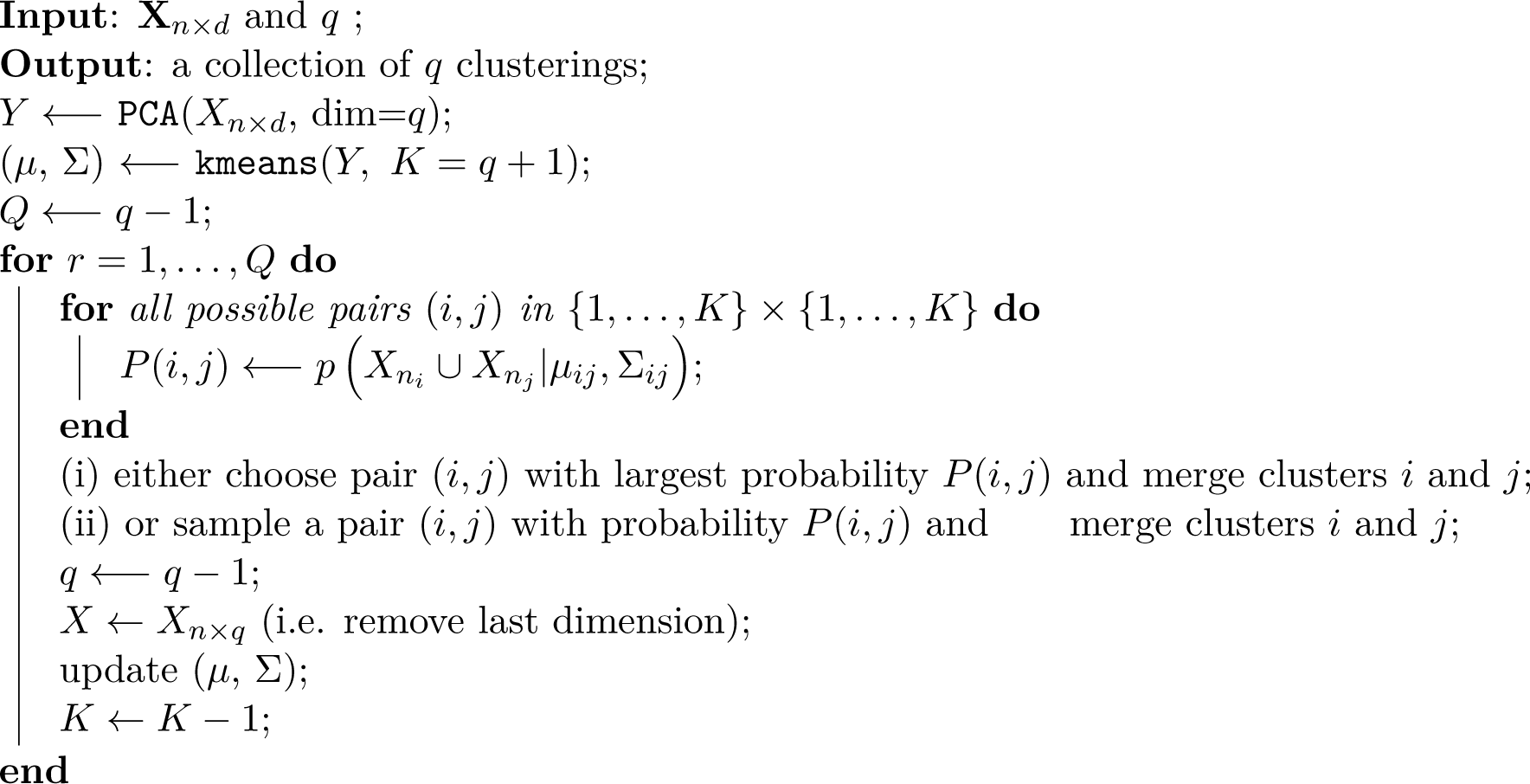

**Algorithm 1:** The *pcaReduce* algorithm. Here **X***_n_×_d_* is a gene expression matrix with *n* cells (given in rows) and *d* genes (in columns); *q* is the number of dimensions – effectively this refers to the number of levels in the hierarchy; *μ_ij_* and Σ*_ij_* definition are given in Equation (1); (i) and (ii) denote two different merging settings.

**Figure 2:**
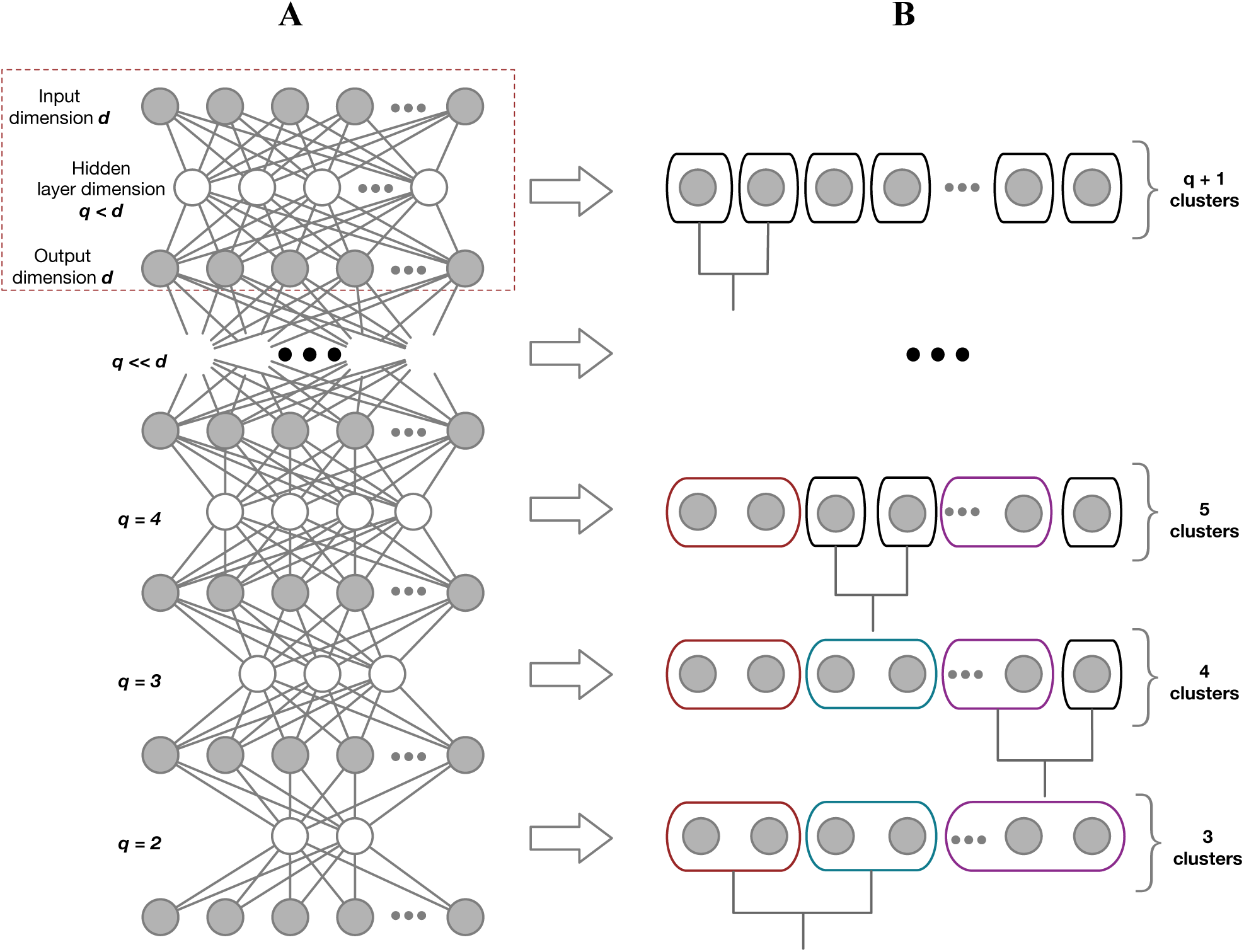
Method illustration using an autoencoder network. Clustering is applied to the data representation at each linear hidden layer. If there are *K −*1 linear hidden units, the data is projected into a subspace spanned by the top *K −*1 principal components. Consistency between the clusterings at each layer is maintained by enforcing a hierarchical constraint. (A) Graphical interpretation of an autoencoder network(s). (B) Corresponding hierarchical structure.

#### Motivation

The algorithm we propose is driven by the expectation that information pertaining to large, broad classes of cell types is likely be contained in the subspace defined by the principal components explaining the most variability in the expression data. Fine-scale cluster sub-structure that is related to more subtle cellular states would be defined by information contained in the principal directions explaining the least total variability.

Therefore, if we cluster initially in a subspace defined by a certain number of principal components, we would expect that if we remove some of those dimensions, the detailed sub-structure disappears as some clusters will become indistinguishable in the lower dimensional subspace. However, rather than re-applying a clustering algorithm afresh each time and leaving a collection of unrelated cluster structures, we select a pair of clusters to merge from the existing structure leading to the formation of a hierarchy. The orthogonality of the principal component subspaces means that there is consistency across the different levels, for this reason clustering at each low-dimensional subspace is embedded within higher dimensional spaces.

An alternative way to interpret our model uses the idea of autoencoder networks (Sperduti, 2013; Baldi, 2012) (see Figure 2A, structure highlighted in red dashed line). An autoencoder is a feed-forward neural network consisting of input, hidden and output nodes. The number of nodes in the input and output layers are equal whilst the number of hidden layers is usually less than the input nodes. As a result, data compression (dimensionality reduction) is induced through the hidden layer, because the information in the input layer is forced into a representation that is encoded by fewer parameters. Our approach takes the compressed data representation from the hidden layer that contains *K* − 1 hidden units and using this for clustering in the *K*-th level of the hierarchical clustering structure (see Figure 2B).

**Figure 3:**
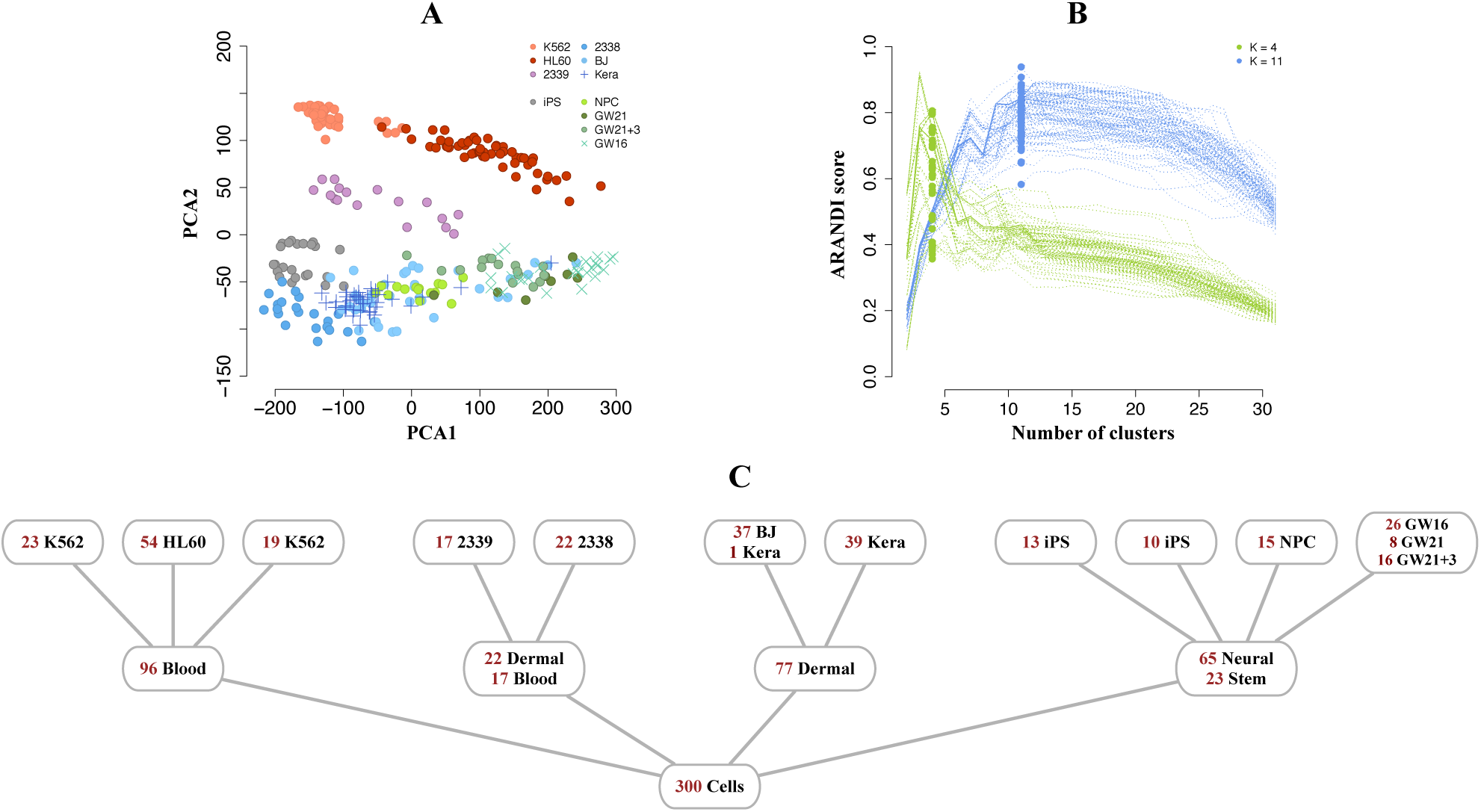
Application of *pcaReduce* to single cell RNA sequencing of 11 cell lines. (A) Projection of the data on to the first two principal components. (B) Performance of *pcaReduce*, the horizontal axis corresponds to a level in the hierarchical cluster structure reported by *pcaReduce*, the vertical axis show the Adjusted Rand Index (ARANDI) score between the tissue-level (green) and cell-line level labels (in blue) and the clustering reported by each level of the hierarchical clustering of *pcaReduce*. Each line correspond to a single run of *pcaReduce* using probabilistic sampling. (C) The most probable cellular hierarchy identified using *pcaReduce*.

Training an autoencoder requires the identification of a set of weight parameters for the hidden and output units to minimise the sum-of-squared reconstruction error between input and output and this is typically done (in general settings) using heuristic approaches (to avoid local minima) combined with backpropagation techniques (gradient descent). In this paper, we only use linear functions for the hidden and output units, thus fitting the autoencoder network is equivalent to performing a principal components analysis (Baldi and Hornik, 1989). However, although not directly examined here, we shall later discuss how this representation leads to further research directions for integrated dimensionality reduction and classification.

## 3 Applications

To demonstrate the performance of *pcaReduce* method, we considered two single cell RNA-seq dataset examples. The first contains a collection of cells originating from diverse biological tissues (Pollen *et al*., 2014); and the second dataset the mouse sensory neuronal cells (Usoskin *et al*., 2015) discussed in the Introduction. These were selected as they contained pre-existing hierarchical cluster structures that can be used to assess unsupervised algorithmic performance. Here we show that *pcaReduce* can be applied to re-capture the known cellular hierarchies and we compare to other popular tools, which are commonly applied to address similar cell sub-typing problems. Below, all examples were implemented using the free statistical computing platform R (www.r-project.org).

### Applications to cells from disparate tissues

We considered a single cell RNA-seq dataset (Pollen *et al*., 2014) that possesses hierarchical complexity. The example contains of 300 cells whose transcriptional measurements are taken across 8,686 genes (see Supplementary Information, Section A for further details on data preparation). The data were derived from 11 cell types: K562 – myeloid (chronic leukemia), HL60 – myeloid (acute leukemia), CRL-2339 – lymphoblastoid; iPS – pluripotent; CRL-2338 – epithelial, BJ – fibroblast (from human foreskin), Kera – foreskin keratinocyte; NPC – neural progenitor cells, GW(16, 21, 21+3) – gestational week (16,21, 21+3 weeks), fetal cortex (see Figure 3A). In addition, as specified in the original study by Pollen *et al*., 2014, these cell types could also be grouped into four disparate tissue sources: blood, stem, skin and neural tissues. We will use the cell lines and tissue-level classifications as ground-truth cellular classes in our performance assessment.

We applied *pcaReduce* to this dataset to construct cellular hierarchies. First, we initially projected the data into the subspace spanned by the first 30 principal components following a PCA and performed an initial *K*-means clustering to get initial cluster assignments (using *K* = 31 clusters) (Ding and He, 2004). After this, we applied different merging strategies to construct the cellular hierarchies: first, when merging is performed based on the most probable cluster merge value (see (i) in Algorithm 1) and, secondly, when merge candidates are probabilistically sampled (see (ii) in Algorithmic overview section). The former gives a single hierarchical clustering whilst the latter can give a range of candidates hierarchies based on repeated sampling.

We compared the clusterings given by *pcaReduce* at level *K* = 4 and *K* = 11 to the *true* tissue and cellular classifications respectively using the Adjusted Rand Index (Hubert and Arabie, 1985). Figure 3B illustrates the performance of *pcaReduce* using the sampling-based merge operation where each line corresponds to a single run of the method. Although, *pcaReduce* has no knowledge of the true number of cell line or tissue labels, the correspondence between the hierarchical clustering output of *pcaReduce* and the true classification peaks at around levels 4 and 11 of the hierarchies respectively, which is reassuring.

In order to gain an understanding of the misclassifications, we looked specifically at the most probable hierarchical structure identified using *pcaReduce* (Figure 3C). Compared to the known cell line and tissue labels (see Supplementary Figure 1A), the 11-cluster structure given by *pcaReduce* did not fully differentiate the 11 cell types. This is not unsurprising since the 11 cell types included a set of three maturing neural cell types (GW16, GW21 and GW21+3) that are highly related. Interestingly, *pcaReduce* grouped these cell types together, which is not an entirely inappropriate operation since the expression variation between the maturing neural cells maybe relatively low compared to unrelated cell types. There was also some class splitting, for example, two sub-groups of K562 cells were identified. Figure 3A qualitatively indicates that this may make sense as some K562 cells were closer in overall expression to HL60 cells than other K562 cells.

At the 4-cluster level the assignments given by *pcaReduce* gave some interesting group structures. The ground truth tissue-level classification assumed the existence of blood, neural, dermal and stem cell types but *pcaReduce* identified that the CRL-2338 and CRL-2339 cell lines should form a group. This is interesting as CRL-2338 is a cell line derived from a primary stage IIA, grade 3 invasive ductal carcinoma and CRL-2339 is a B lymphoblastoid cell line initiated from peripheral blood lymphocytes from the same patient. Pluripotent stem cells (iPS) were also grouped by *pcaReduce* with neural progenitor cells (NPC). Overall, whilst *pcaReduce* did not give a 4-cluster classification that was identical to the original tissue classifications, the output produced are not necessarily nonsensical. In comparison, the output of standard hierarchical clustering shows very little obvious structure (see Supplementary Figure 1/2).

**Table 1:**
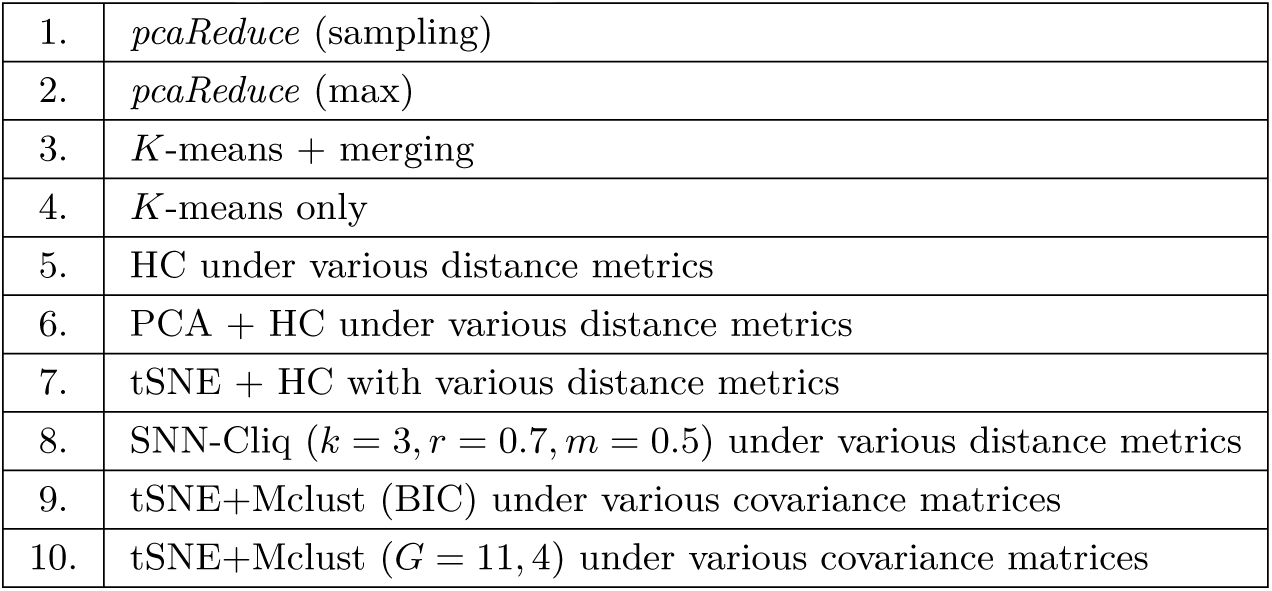
Methods used in performance comparison. Further details regarding parameter setup and running specifications are given in the Supplementary Information, Section A.

**Figure 4:**
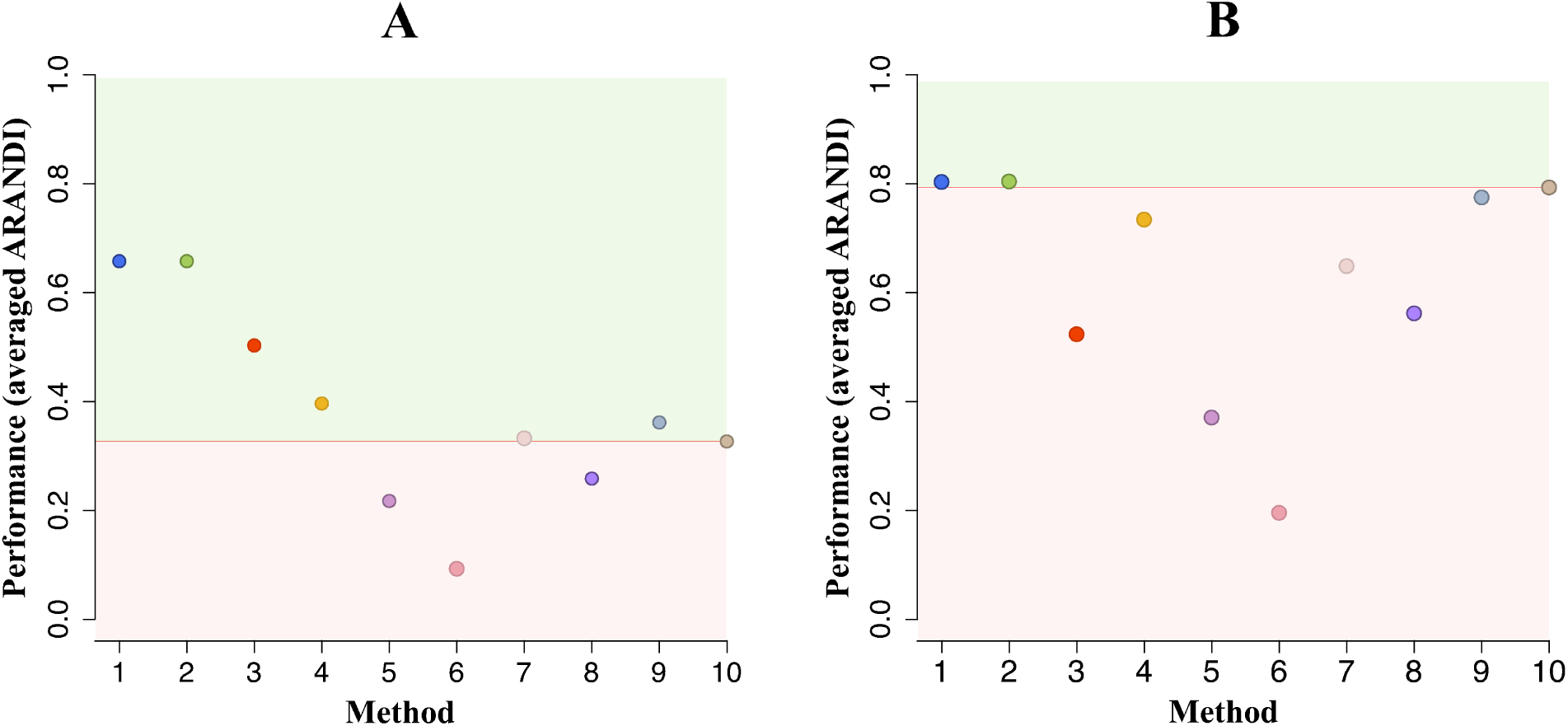
Performance comparison on cell line data. Classification performance against known tissue-level and (B) cell-line level labels. Points positioned above red line (in green background) indicate better performance, whereas points below (in red background) illustrate poorer performance relative to the benchmark (Method 10). Numbers 1 *−* 10 correspond to clustering methods in Table 3.

In order to fully assess the performance of *pcaReduce*, we compared it to a set of alternative approaches (see Table 3 for a full list), which are often applied to address similar, cell type discovery, problems. This includes popular methods such as: *K*-means, hierarchical clustering (HC), Mclust – mixture modelling for model-based clustering (Fraley and Raftery, 2004) combined with tSNE – a nonlinear dimensionality reduction/visualisation technique (Van der Maaten and Hinton, 2008); and a recent single cell method – SNN-Cliq, which determines similarities between cells based on shared nearest neighbours algorithm and performs single cell clustering using graph-theoretical concepts (Xu and Su, 2015). Details regarding all parameters and running specifications for each clustering approach are summarised in Supplementary Information, Section A.

Figure 4 shows the relative performance of different approaches. We used the score of Mclust/tSNE (Method 10) as a benchmark as this was the best performing method for classification of the 11 cell lines. However, whilst some methods had comparable performance to *pcaReduce* in terms of capturing the 11 cell types, their performance diminished for the four cluster tissue-level classifications. Overall, *pcaReduce* was the only method that could provide strong clustering results for both the cell line and tissue classifications. Interestingly, the *K*-means with merging algorithm (Method 3), which can be interpreted as a simpler variant of *pcaReduce* that does not involve dimensionality reduction steps, performed relatively poorly. A more detailed breakdown of the performance comparison is given in Supplementary Figure 3. This shows the performance sensitivity of standard hierarchical clustering algorithms to the choice of distance metric and the dependency on initialisation for the outcome of the *K*-means algorithm.

### Applications to mouse neuronal cells

We next returned to the mouse neuronal cell dataset discussed in the Introduction that contains measurements across 25,334 genes (Usoskin *et al*., 2015). The study classified cells according to four principle neuronal groups: non-peptidergic nociceptor cells (NP), peptidergic nociceptor cells (PEP), neurofilament containing cells (NF), and tyrosine hydroxylase containing cells (TH) (Figure 5A). In addition to this, it was suggested that the NP, PEP and NF cells possessed further subtypes (Figure 1B,C). We now examined whether *pcaReduce* could recover this three layer classifications within its hierarchical output without the use of marker genes.

We applied *pcaReduce* and computed the correspondence between the 4-, 8- and 11-cluster structures and those in the original study. Figure 5B shows that the absolute classification accuracy was relatively low compared to the previous cell line experiment. This is unsurprising as the four predominant neural cell groups form a complex cluster pattern in the subspace spanned by PC2-4 (see Figure 5A) and would be hard to segregate in an entirely unsupervised way as we propose. The most probable structure given by *pcaReduce* identifies three groups that are predominantly dominated by NP, TH and NF cells respectively (Figure 5C). However, these groups contains some contamination due to the fact that the clusters overlap and there is a miscellaneous fourth group that contains most of the PEP cells (49/63) but is actually dominated by NP cells. The application of tSNE – a non-linear dimensionality reduction technique – did not well-separate the four groups (Supplementary Figure 7) and it would not be obvious, without known markers, how to delineate each group.

Figure 6 shows that the performance of *pcaReduce* was consistently amongst the best performing relative to other methods. Note that although the benchmark algorithm (Method 10) provides the best performance for each level of clustering, it should be noted that each set of cluster assignments produced are not hierarchically linked as in *pcaReduce* and so they maybe inconsistent. A more detailed breakdown of the performance results is given in Supplementary Figure 5.

Here we have evaluated all methods, including *pcaReduce*, on a full dataset without prior filtering of any cell/gene subsets. For this reason the identification of cell subtypes proves to be particularly challenging, since the neuronal sub-classes (i.e. *K* = 8, 11) appear to be poorly separable without prior knowledge of cellular biomarkers. This is especially evident from PCA plots summarised in Supplementary Figure 6, where we plot pairwise combinations of various principle components and highlight cells that should correspond to neuronal subtypes: NF3 and NF4 (lower-left) and NP2 and NP3 (upper-right). In addition, we highlight the cell subtype separability issue using non-linear dimensionality reduction/visualisation technique, tSNE (see Supplementary Figure 7B-C), that suggests (at least visually) alternative cell assignments into subclasses.

**Figure 5:**
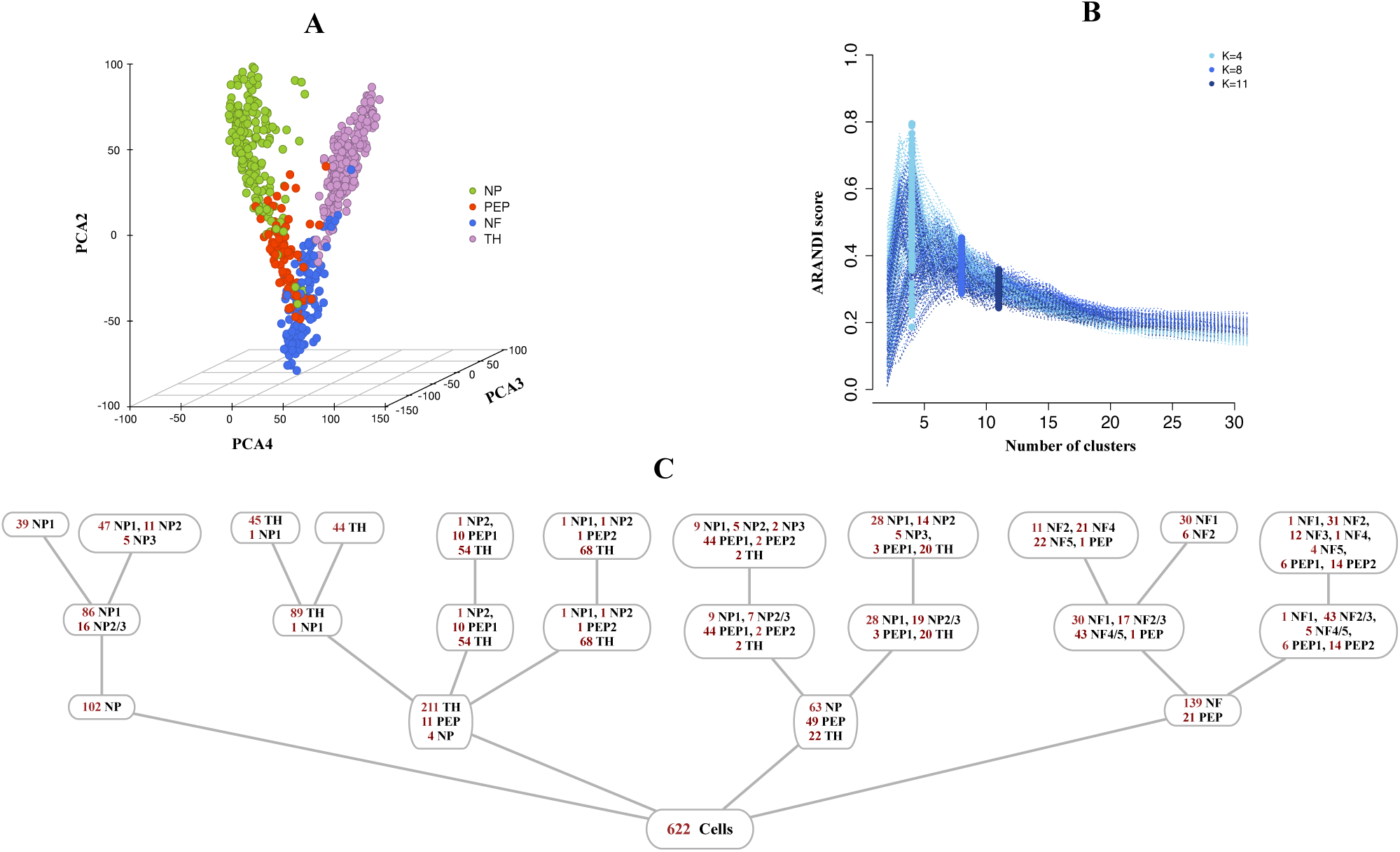
Application to single cell mouse neuronal data. (A) Data projected on to PC2-4 for visualisation and coloured by the four major neuronal cell types. (B) Clustering performance of *pcaReduce*. (C) Cellular hierarchy identified using *pcaReduce*.

**Figure 6:**
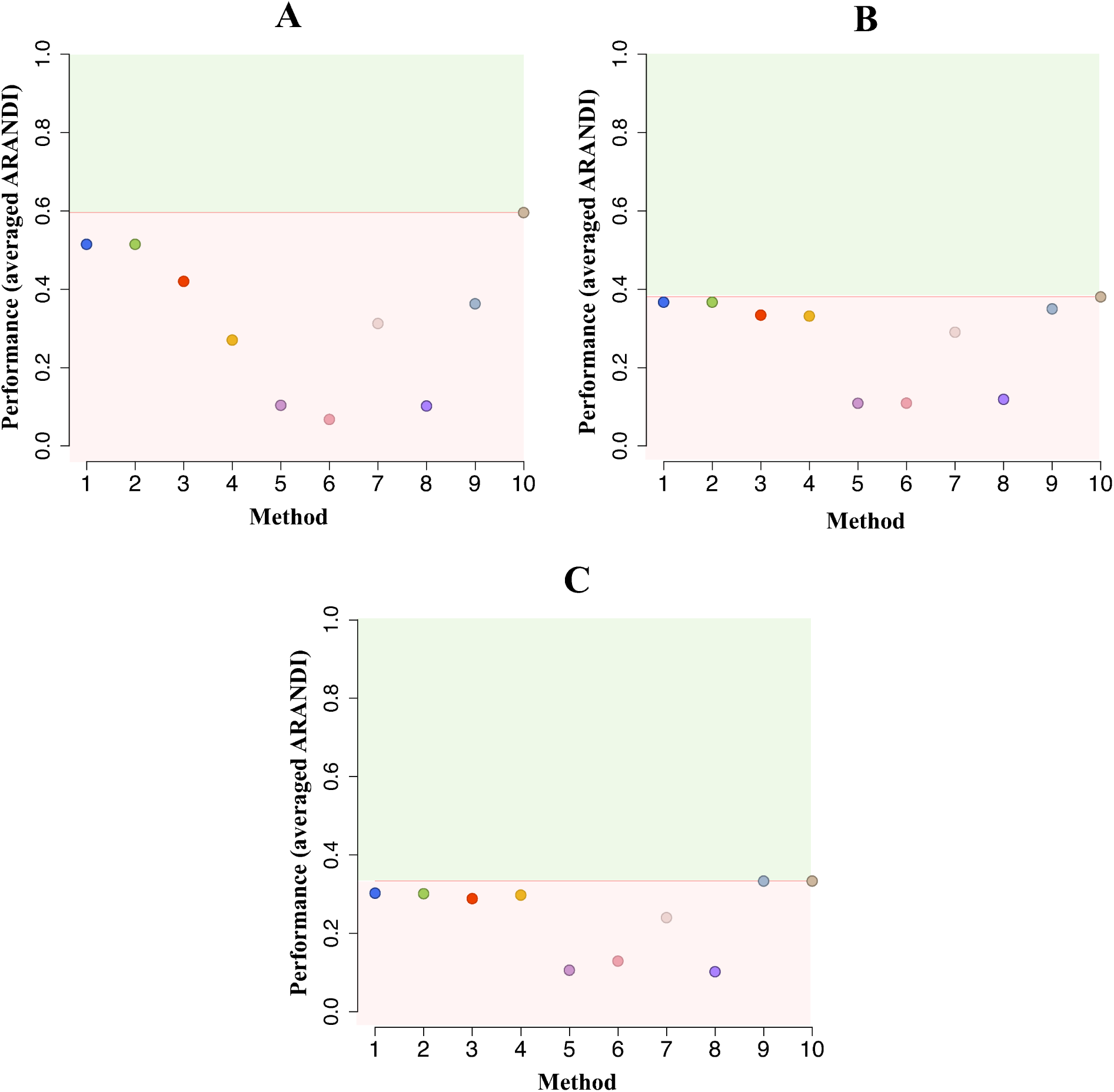
Performance comparison on mouse neuronal data. Classification performance against known (A) the four major neuronal classes and (B, C) further subtypes identified in the original study (Usoskin *et al*., 2015). Points positioned above red line (in green background) indicate better performance, whereas points below (in red background) illustrate poorer performance relative to the benchmark (Method 10). Numbers 1 − 10 correspond to clustering methods in Table 3.

## 4 Discussion and Conclusions

In this paper we have presented an unsupervised hierarchical clustering approach for the identification of putative cell sub-populations from single-cell transcriptomics profiles. Clustering occurs in a linearly transformed subspace obtained from principal component directions and, at each level of our hierarchical clustering structure, the similarity between clusters is measured in subspaces of *decreasing* dimensionality by discarding principal directions as the number of clusters decreases. In doing so, we presume that the variation contained in the first principal components corresponds to the features of broad cellular classes, whilst fine-scale variation in lower principal directions correspond to the features of detailed cellular sub-structure.

We applied this technique to two illustrative single cell datasets from the recent literature and showed that, compared to a variety of existing clustering tools, our approach was able to better recapitulate pre-existing cluster structures across both – broad *and* detailed cellular states; further, this was achieved *simultaneously* in a hierarchical fashion. However, we note that the absolute performance of ours and other techniques can be rather limited in an unsupervised setting, and further research is required to combine local and global feature selection alongside clustering/classification techniques is necessary in order to better identify real cell types and states.

Our clustering approach can be interpreted within an autoencoder network framework. This offers the possibility for further method extensions using non-linear transformations for dimensionality reduction or data compression. We are currently exploring this, and the application of non-hierarchical, multi-label assignment algorithms for cell state classification problems where a strict hierarchical structure is not appropriate.

## Acknowledgement

The authors are supported by a Wellcome Trust Core Award Grant Number 090532/Z/09/Z, the Li Ka Shing Foundation Oxford-Stanford Big Data for Human Health Seed Grant and a UK Medical Research Council New Investigator Research Grant (Ref. No. MR/L001411/1).

## A Further methods

### Note on low level single cell data processing and gene expression matrix preparation

We have downloaded RNA-seq dataset from NCBI repository (http://www.ncbi.nlm.nih.gov) under accession number SRP041736, which contains transcriptional profiles of 347 singles cells. Next, we have converted the Sequence Read Archive (SRA) files into *fastq* files using SRA Toolkit (http://www.ncbi.nlm.nih.gov/sra/), and used TopHat-[5] to perform genomic mapping of pair-end reads to the latest human reference genome GRCH38 (http://www.ensembl.org/info/data/ftp/index.html). We have used R package called *Rsubread* [2] to assign mapped sequencing reads to genomic features, i.e. to perform transcript counting, which was achieved using function *featureCounts*.

In order to construct a gene expression matrix for higher level analysis, we have performed basic cell and gene filtering: (a) From 347 samples we have focused on a subset of 301 cells (a subset without ERCC validation cells and bulk sample), which were used in the main study by Pollen et al. [3]. In addition, we have removed one cell that had 0 expression levels across all genes, this left us with a 300 cells in total. (b) For gene filtering we have used R package called *edgeR* [4], we kept those genes that fulfilled at least 100 counts per million (cpm) in at least 10 samples; this left us with 8686 genes in total. Lastly, we have transformed gene expression counts to a logarithmic *ij ij* values; more precisely, values *x*_*ij*_ in matrix *X* were obtained by 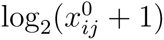 where 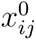 are read counts of a gene.

### Note on clustering tool comparison

- **K-means and merging**. Using PCA we project initial dataset to *q* = 30 and partition it into *K* = 31 clusters using K-means. Next, we merge two clusters together using sampling as a merging criterion. However, in the next step instead of dropping off the last dimension (i.e. instead of *q ← q −*1 in Algorithm 1 in main paper), we keep *q* = 30 fixed, and continue merging clusters as described before. We repeat these steps 100 times. This small method alteration illustrates the importance of gradual dimensionality reduction, as it affects the *pcaReduce* performance quality on both datasets (see Method 3, averaged ARANDI scores in Figures 4 and 6 in main paper).
- **K-means only**. We ran the K-means clustering algorithm for a full dataset 100 times without prior dimensionality reduction step; using either the true number of clusters, e.g. for the Pollen data set *K* = 4 and *K* = 11, and computed the ARANDI scores between these clusterings and known cellular labels (e.g. see Method 4 in Figure 4 A and B respectively).
- **Hierarchical clustering**. We ran the hierarchical clustering algorithm with all possible distance measures: Euclidean, maximum, Manhattan, Canberra, and binary. Each time we cut hierarchical tree at e.g. *K* =4 and *K* = 11 and compute ARANDI scores between these clusterings and known cellular labels (e.g. see Method 5 in Figure 4 A and B respectively).
- **PCA followed by Hierarchical clustering**. Using PCA we project initial dataset to *q* = 30 and run hierarchical clustering algorithm with all possible distance measures. Each time we cut hierarchical tree at e.g. *K* = 4 and *K* = 11 and compute ARANDI scores between these clusterings and known cellular labels (e.g. see Method 6 in Figure 4 A and B respectively).
- **RtSNE followed by Hierarchical clustering**. Using RtSNE we project data on to two-dimensional space, we select the number of dimensions that should be retained in the initial PCA step to be 30 (the same as *q* = 30), and set perplexity parameter to be 30, further we use accuracy parameter to be default, *θ* = 0.5. We ran the hierarchical clustering algorithm with a range of possible distance measures: Euclidean, maximum, Manhattan, Canberra, and binary. Each time we cut hierarchical tree at e.g. *K* = 4 and *K* = 11 and compute ARANDI scores between these clusterings and known cellular labels (e.g. see Method 7 in Figure 4 A and B respectively).
- **SNN-Cliq**. We ran the SNN-Cliq [8] tool on a full dataset using a range of possible distance measures: Euclidean, maximum, Manhattan, Canberra, and binary. We keep default settings, i.e. *k* = 3, which is the size of the nearest neighbours, *r* = 0.7, which is a parameter for finding quasi-cliques, and *m* = 0.5 – a merging parameter; the later two parameters affects cluster compositions. Next, we compute ARANDI score between true clusterings, *K* = 4 and *K* = 11, and clusterings determined by SNN-Cliq. (e.g. see Method 8 in Figure Figure 4 A and B respectively). It is worth noting that by examining various parameter *k, r, m* combinations, one potentially could achieve a greater agreement between estimated and known clusterings. However, this could also pose some difficulties in situations where cluster labels are unknown.
- **RtSNE in conjunction with Mclust**. We use t-SNE [7] (R package *Rtsne* [6]) to project dataset on to two-dimensional space, we select the number of dimensions that should be retained in the initial PCA step to be 30 (the same as *q* = 30), and set perplexity parameter to be 30, further we use accuracy parameter to be default, *θ* = 0.5. Next we use model based clustering, Mclust [1], with all possible covariance models (for further description see Supplementary Figure 3) to cluster projected dataset. We use the following strategies:
  1. To determine the number clusters that best describes provided dataset, we use Bayesian information criterion (BIC). We test the number of mixture components in the range of *G* = 1 : 31. Next we compute ARANDI scores between the true cluster labels, *K* =4 and *K* = 11, and labels identified with BIC (e.g. see Method 9 in Figure 4 A and B respectively).
  2. Alternatively, using Mclust we fit two finite mixture models with fixed number of mixture components, *G* = 4 and *G* = 11. Again we compute ARANDI scores between true clusterings and clusterings identified by Mclust (e.g. see method 10 in Figure 4 A and B respectively).

## B Applications to cells from disparate tissues

**Figure 1:**
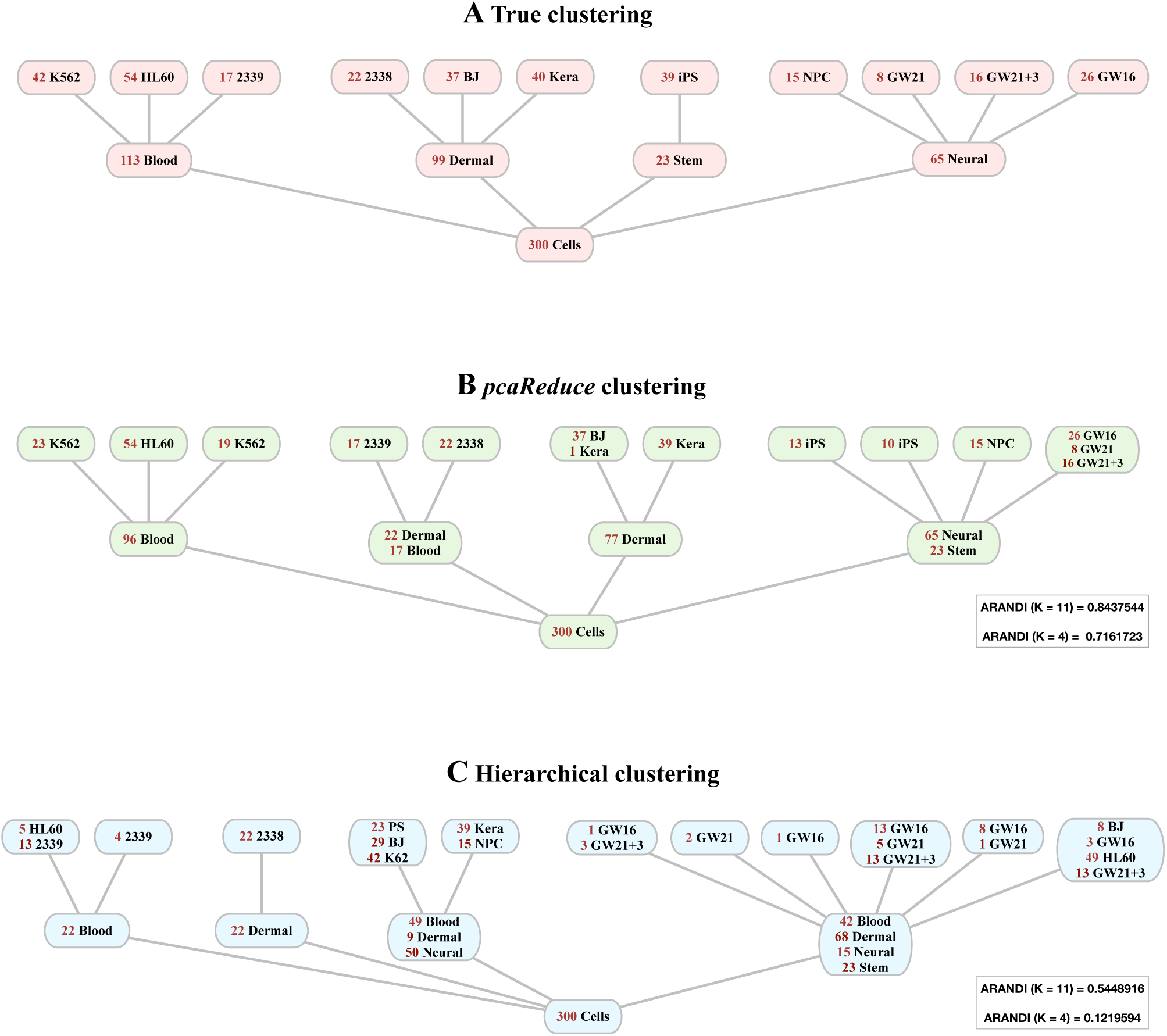
Application to cells from disparate tissues. (A) Illustrate hierarchical relationships between cells based on known cellular labels. (B) Hierarchical relationships between cells based on cellular labels identified by single run of *pcaReduce* under max merging criterion. Also ARANDI scores between true clusterings, *K* = 4 and *K* = 11, and identified clusterings by *pcaReduce* are summarised in grey rectangle. (C) Hierarchical relationships between cells based on cellular labels identified by hierarchical clustering (HC) algorithm; ARANDI scores between true clusterings, *K* = 4 and *K* = 11, and identified clusterings by HC are summarised in grey rectangle.

**Figure 2:**
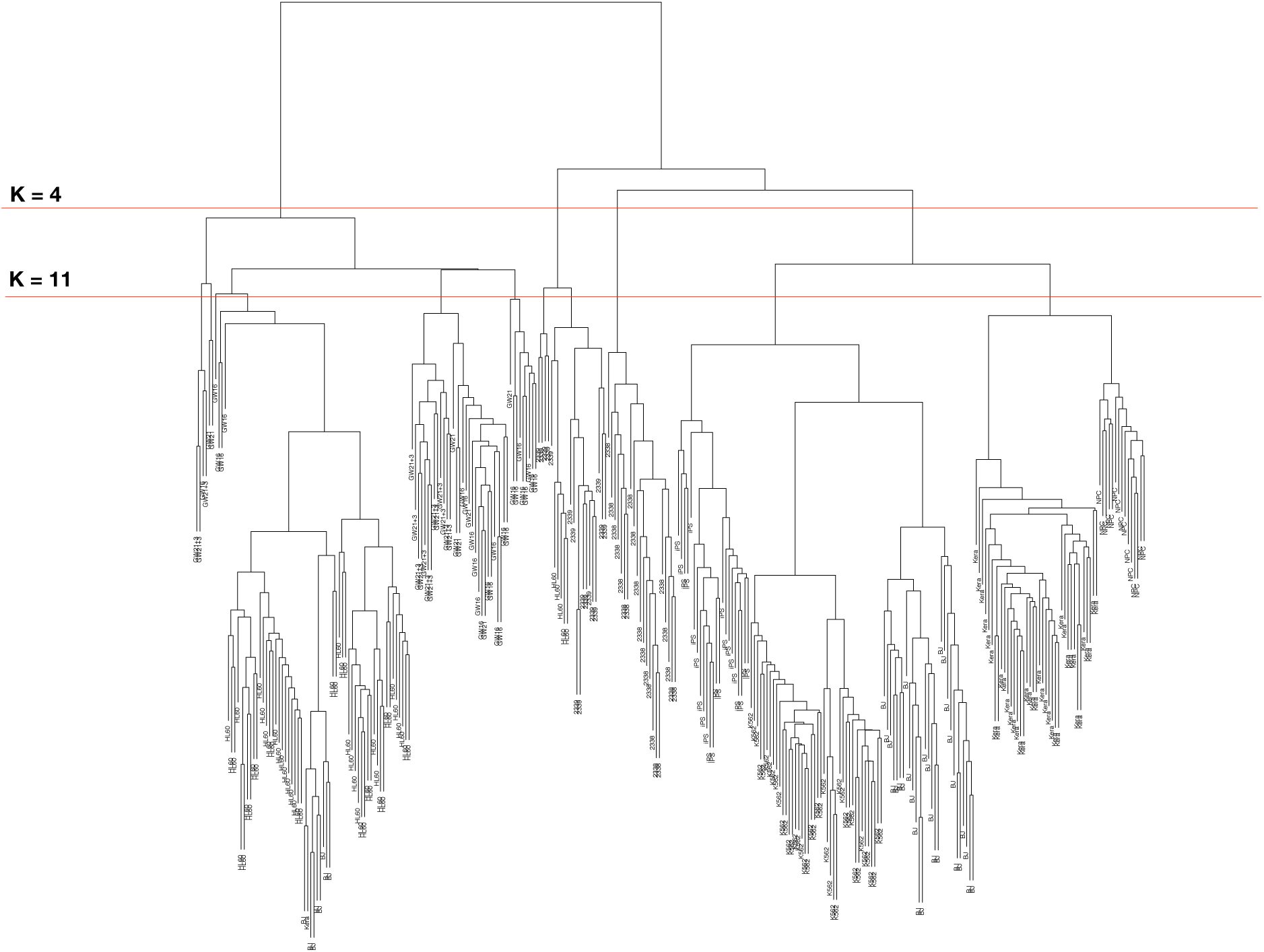
Application to cells from disparate tissues. Illustrates a tree structure obtained using hierarchical clustering. Red guide lines illustrate clusterings obtained by cutting tree at different level: K = 4 and K = 11.

**Figure 3:**
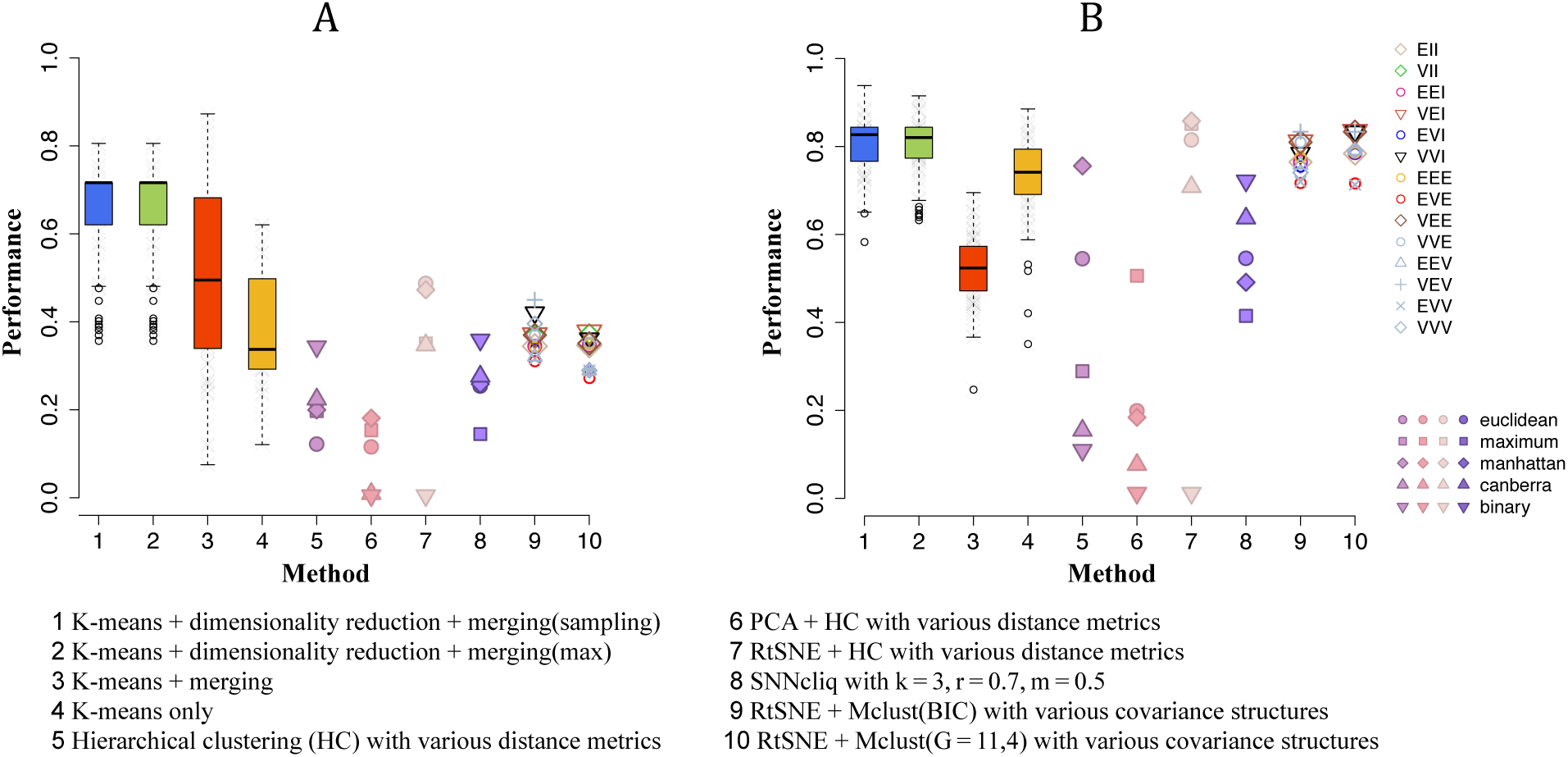
Application to cells from disparate tissues. Boxplots 1–4 are based on 100 runs of each method; methods 5–8 are evaluated based on all possible distance metrics, and 9–10 based on all possible covariance structures. (A) ARANDI score between true clustering, *K* = 4, and clusterings identified by 1–10 approach. (B) ARANDI score between true clustering, *K* = 11, and clusterings identified by 1–10 approach.

## C Applications to mouse neuronal cells

**Figure 4:**
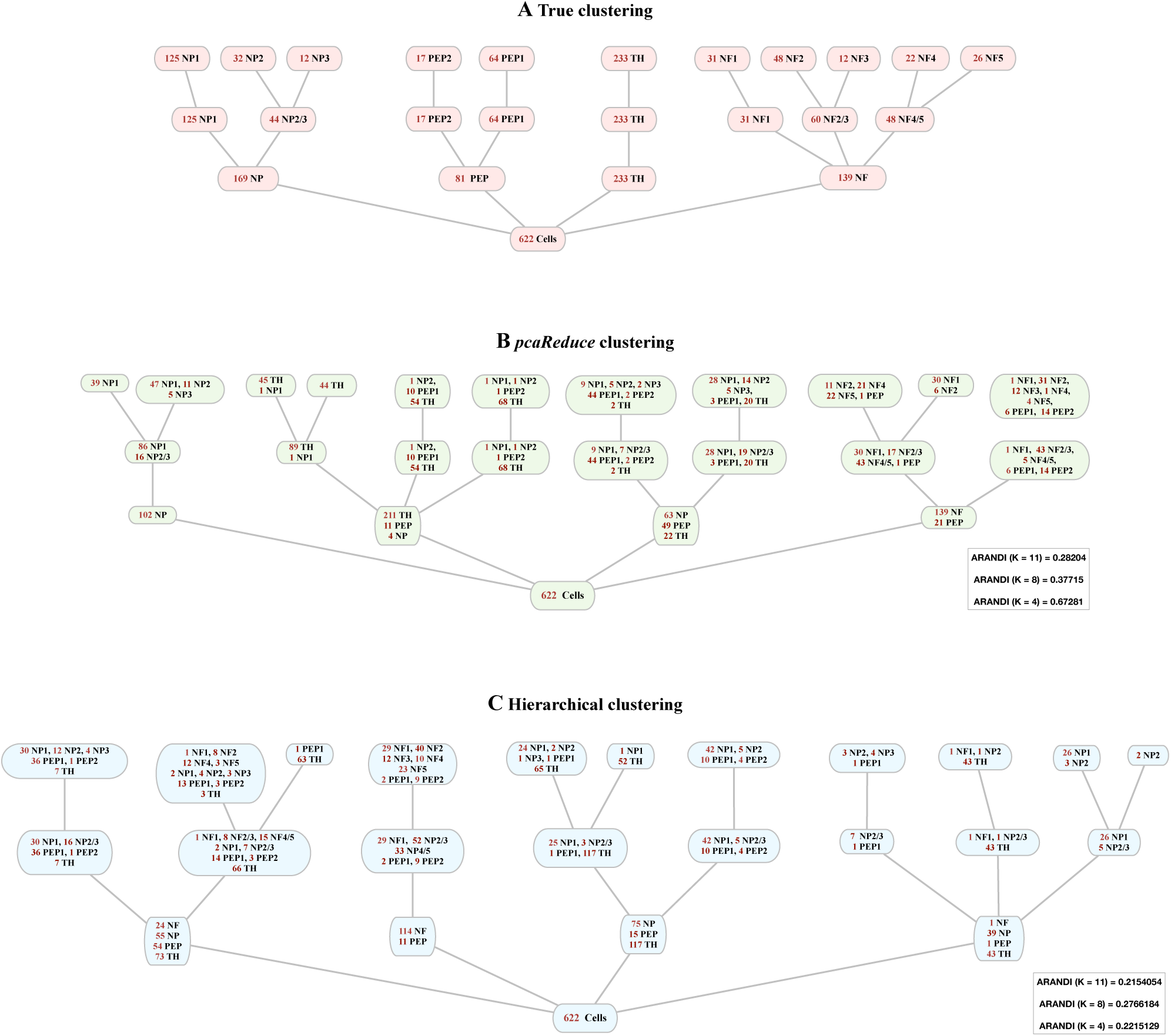
Applications to mouse neuronal cells. (A) Illustrate hierarchical relationships between cells based on known cellular labels. (B) Hierarchical relationships between cells based on cellular labels identified by single run of *pcaReduce* under max merging criterion. Also ARANDI scores between true clusterings, *K* = 4, 8, 11, and identified clusterings by *pcaReduce* are summarised in grey rectangle. (C) Hierarchical relationships between cells based on cellular labels identified by hierarchical clustering (HC) algorithm; ARANDI scores between true clusterings, *K* = 4, 8, 11, and identified clusterings by HC are summarised in grey rectangle.

**Figure 5:**
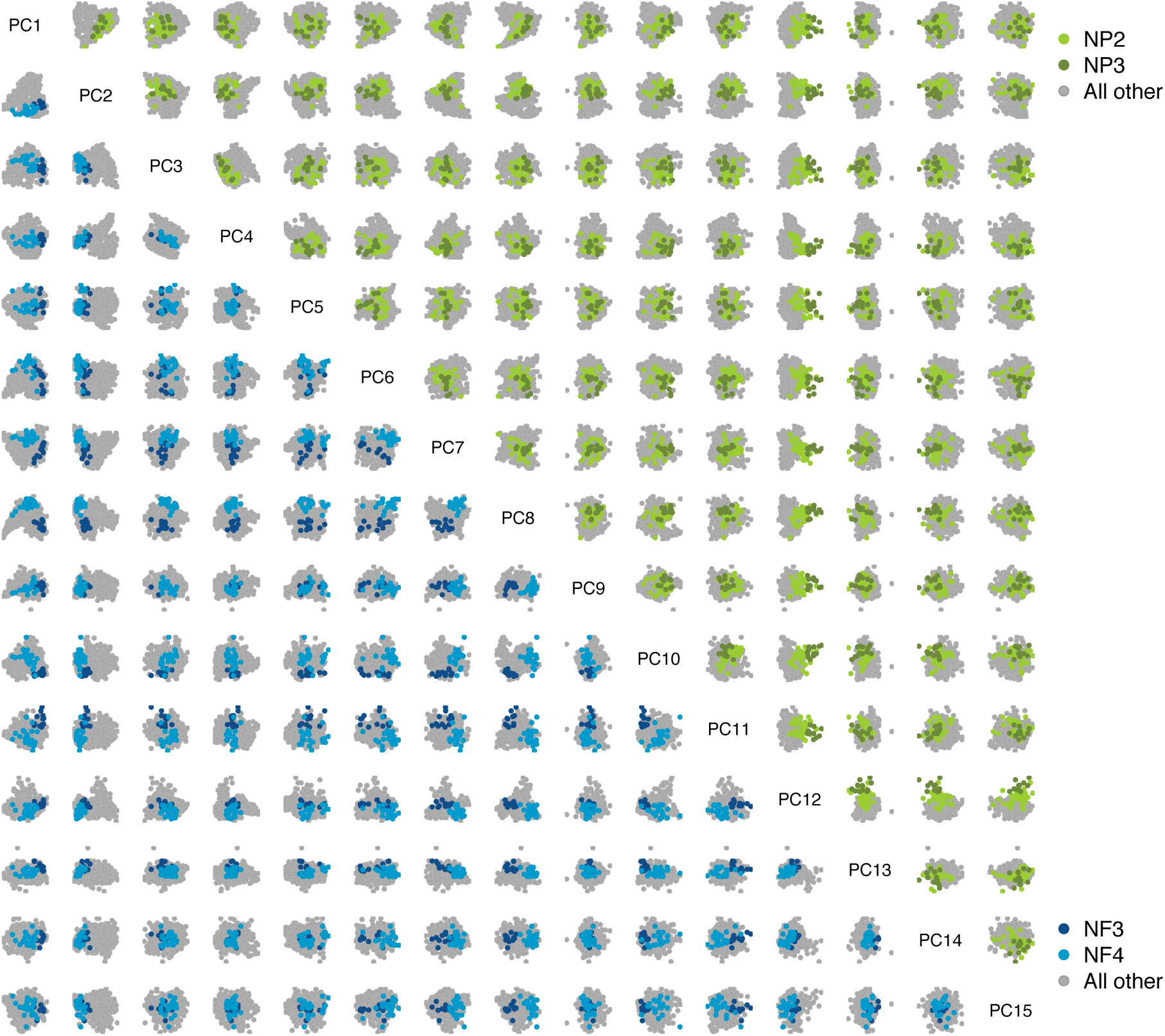
Applications to mouse neuronal cells. Scatterplots illustrate the first 15 principle directions. In left triangle NF3 and NF4 cells are highlighted in blue and dark blue points. In right triangle NP2 and NP3 cells are highlighted in green and dark green points.

**Figure 6:**
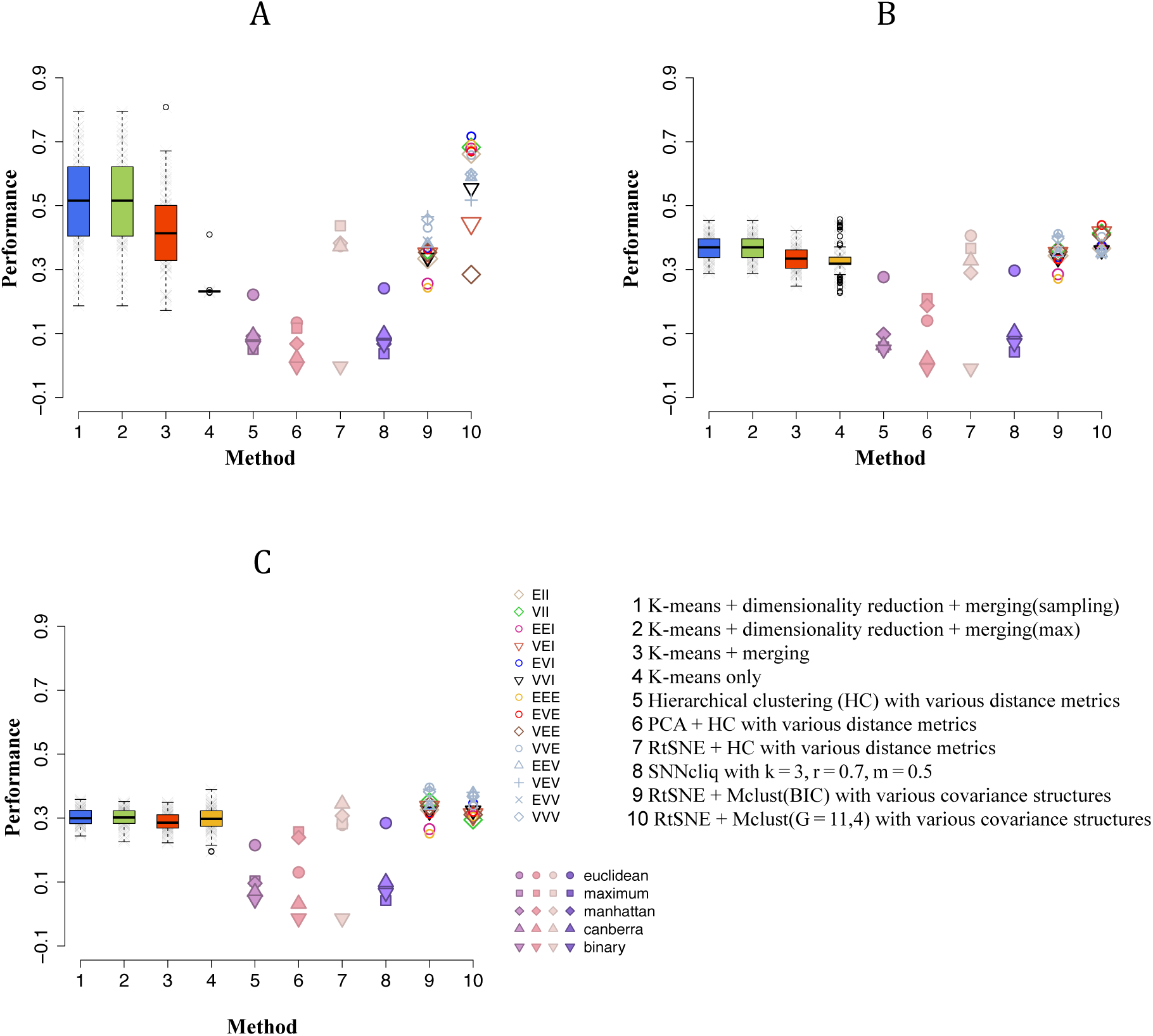
Applications to mouse neuronal cells. Boxplots 1–4 are based on 100 runs of each method; methods 5–8 are evaluated based on all possible distance metrics, and 9–10 based on all possible covariance structures. (A) ARANDI score between true clustering, *K* = 4, and clusterings identified by 1–10 approach. (B) ARANDI score between true clustering, *K* = 8, and clusterings identified by 1–10 approach. (C) ARANDI score between true clustering, *K* = 11, and clusterings identified by 1–10 approach.

**Figure 7:**
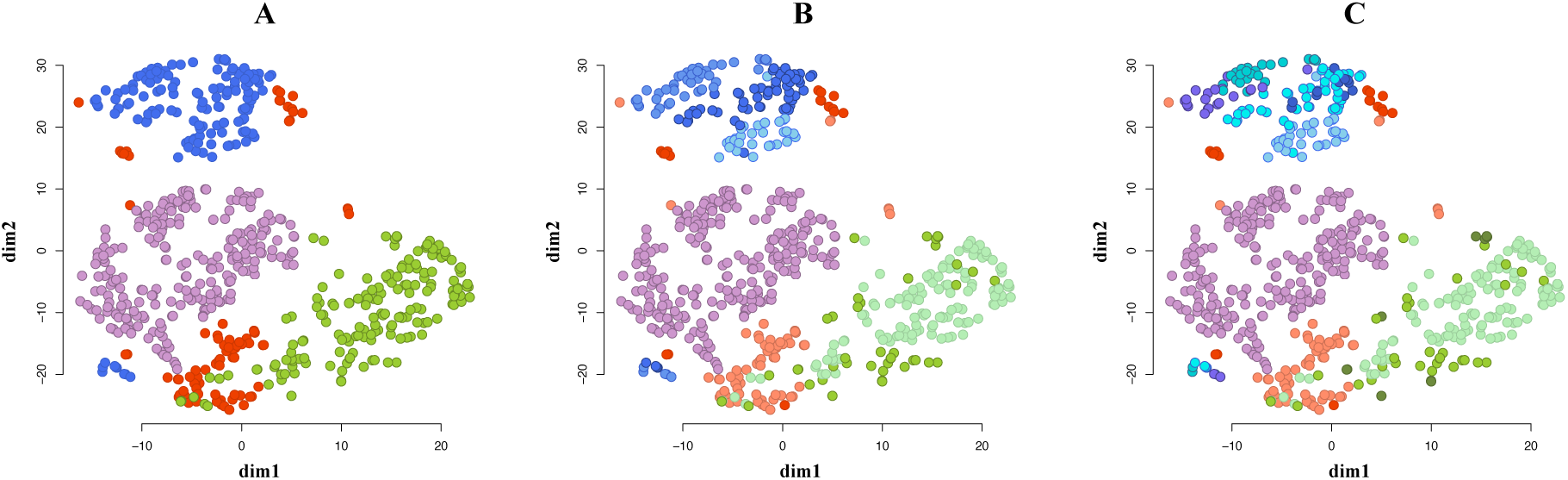
Applications to mouse neuronal cells. All plots illustrate the same dataset using RtSNE visualisation tool applied for projecting data on to two-dimensional space, here cells are coloured based on available labels and illustrate various clustering patterns: (A) 4, (B) 8, and (C) 11 clusters.

